# Genetic basis of Cassava (*Manihot esculenta* Crantz) plant architecture and its relevance for selection of farmer-preferred varieties

**DOI:** 10.64898/2026.02.11.705251

**Authors:** Pamelas M. Okoma, Siraj S. Kayondo, Ismail Y. Rabbi, Chinedozi Amaefula, Luciano Rogerio Braatz de Andrade, Lydia C. Jiwuba, Joseph Onyeka, Chiedozie Egesi, Jean-Luc Jannink, The NextGen Cassava Consortium

## Abstract

Plant architecture, the spatial configuration of stems, branches, leaves, and inflorescences underpins essential physiological functions such as light capture, assimilate partitioning, flowering, and ultimately, yield. In cassava (*Manihot esculenta*), architectural traits such us plant height, branching level, and plant shape are agronomically important yet remain underexploited in breeding. Here, a large-scale analysis was conducted using phenotypic and genomic data from more than 14,000 cassava accessions evaluated across 34 field locations in Nigeria between 2010 and 2021, encompassing the national breeding programs of the National Root Crops Research Institute and the International Institute of Tropical Agriculture. The study aimed to dissect the genetic architecture, environmental stability, and breeding relevance of four key traits: plant full height, height to first branching, the branching level number (BranchlevelNum) and plant shape. Phenotypic analyses across breeding stages revealed consistent variation in plant height, branching height, and branching intensity, reflecting the cumulative effects of selection and evaluation across environments. Broad-sense heritability estimates ranged from 0.41 to 0.72, with BranchlevelNum and Cylindrical shape exhibiting strong genetic control and weak correlations with yield components, indicating their suitability for independent improvement. Genome-wide association analyses identified significant loci associated with BranchlevelNum, including a major region on chromosome 2 and an additional locus on chromosome 13, collectively explaining approximately 11% of the phenotypic variance. Candidate genes within these regions included regulators of meristem activity and hormone-related pathways, supporting a developmental basis for branching variation. Genomic prediction accuracy for BranchlevelNum reached 0.44, comparable to values reported for key agronomic traits in cassava.

These results demonstrate that branching-related architectural traits are genetically tractable, largely independent of yield, and amenable to genomic selection. The findings support the integration of BranchlevelNum and plant shape into ideotype-driven breeding frameworks aimed at improving flowering efficiency, canopy structure, and field performance in cassava.

**Author Summary:** Cassava is a major food crop, and its plant shape plays an important role in how easily it can be grown, harvested, and improved through breeding. Traits such as plant height, branching, and canopy form affect flowering, seed production, and field management, yet they have received much less attention than yield or disease resistance.

In this study, we examined plant architecture using field and genetic data from more than 14,000 cassava plants grown across Nigeria over twelve years. We focused on key traits describing plant height, branching level, and overall plant shape. We found that branching level is strongly controlled by genetics, remains stable across environments, and can be predicted accurately using genomic data. We also identified specific regions of the cassava genome linked to branching behavior.

Our findings show that plant architecture can be improved using modern breeding tools without compromising yield. Incorporating branching traits into breeding programs can help develop cassava varieties that flower more reliably and perform better in farmers’ fields.

## Introduction

The comprehensive three-dimensional organization of plant structure referred to as plant architecture (PArt) is defined by the type, number, and spatial arrangement of organs such as stems, branches, leaves, and inflorescences. This architectural blueprint includes stem height, branching pattern, canopy form, and leaf phyllotaxy, all expressed within a spatiotemporal environmental context. PArt is predominantly shaped by developmental processes governing shoot branching, stem elongation, and inflorescence morphology [1–5]. In turn, the resulting plant architecture influences critical physiological functions, including light capture, source–sink dynamics, reproductive competence, and ultimately, crop productivity [2,6]. Consequently, architecture has considerable agronomic and ecological importance, affecting resource-use efficiency, adaptation to climate stressors, and suitability for mechanized cultivation. Although architectural traits are under genetic control, they remain developmentally plastic and environmentally responsive. Their complexity arises from interactions among genetic regulators, hormonal pathways (notably auxin, cytokinin, and strigolactones), and environmental cues such as temperature, light, and nutrient availability [7,8]. This plasticity, while beneficial for adaptation, complicates selection in breeding programs that aim for structural consistency across variable field environments.

In cassava (*Manihot esculenta* Crantz), a clonally propagated staple in sub-Saharan Africa, plant architecture is a trait of growing importance. Beyond its relevance to photosynthetic efficiency and reproductive development, cassava architecture strongly influences mechanization potential, harvesting ease, and propagation efficiency. Farmers typically favor erect, non-branching plants, which facilitate field management and optimize stem use for replanting [9]. However, the breeding implications of architecture are complex: early and profuse branching often promotes floral development, which is critical for hybridization, but these traits may yield undesired shapes or low stem vigor [10,11].

Cassava breeders face the challenge of reconciling flowering optimization which benefits from early branching with farmer-preferred ideotypes, which often feature delayed or absent branching. The timing of first branching is tightly linked to the onset of flowering, with delayed floral induction typically associated with increased first branch height and overall plant height [12,13]. While various agronomic interventions such as pruning, photoperiod adjustment, and plant growth regulators can stimulate flowering [14,15], the resulting architectural shifts may not align with ideotype preferences. This trade-off underscores the need for improved understanding of the genetics of cassava PArt.

Although recent genetic advances have accelerated selection for traits such as yield, disease resistance, and root quality [16–18], architectural traits have received comparatively limited attention. Breeding pipelines have incorporated morphological descriptors such as Shape of Plant and Height to First Branching [9,19], yet a comprehensive dissection of the heritable basis and genomic architecture of PArt remains lacking. Meanwhile, multi-environment trials have revealed significant variation in architectural traits such as plant height, branching height, and branching level number, with important implications for ideotype suitability and reproductive development [12,20].

Recent evidence supports the use of architectural traits as indirect selection targets for flowering. For example, BranchlevelNum correlates positively with floral abundance and synchrony [12], and is among the most heritable architecture traits across environments [21]. However, early-branching types may carry undesirable shapes or canopy forms, prompting a need to disentangle trait correlations and identify candidate loci that could enable more precise selection to change one trait while leaving another constant [22,23]. While GWAS studies have identified QTLs for flowering and node number [12], there is a critical knowledge gap regarding plant shape traits such as compactness, openness, or cylindrical form which may better capture farmer shape preferences.

To address this gap, the present study leverages a large multi-environment dataset comprising more than 14,000 cassava accessions evaluated over a decade of field trials across 34 locations in Nigeria. Accessions were drawn from two major breeding programs the International Institute of Tropical Agriculture (IITA) and the National Root Crops Research Institute (NRCRI) and evaluated at successive stages of genetic advancement. We focus on four architectural traits: PlantHeight, FirstBranchHeight, BranchlevelNum, and plant shape (classified as umbrella, compact, open, cylindrical). Our specific objectives were to:

– quantify genetic variance and heritability of key architectural traits across trials and environments;
– assess phenotypic trends across breeding stages and programs;
– examine correlations between architectural and yield-related traits;
– identify genomic regions and candidate genes associated with architectural traits via GWAS;
– evaluate genomic prediction accuracy to inform breeding for architectural ideotypes.

Together, these analyses aim to deepen our understanding of the genetic and phenotypic basis of cassava plant architecture, and to support more informed ideotype development in breeding pipelines that increasingly integrate flowering, structure, and farmer preferences.

## Material and methods

### 1 Field experiments and phenotyping

Phenotypic data was collected on 14,416 accessions from 2 breeding programs, the International Institute of Tropical Agriculture (IITA) and the National Root Crops Research Institute (NRCRI), in Nigeria. Accessions were evaluated at successive stages of selection including clonal evaluation trials (CET), preliminary yield trials (PYT), advanced yield trials (AYT) and uniform yield trials (UYT). A total of 352 trials established under 4 designs (augmented, alpha, completely randomized design and the randomized complete block design) were carried out from 2010 to 2021 across 34 locations in Nigeria. Planting and trial management adhered to technical recommendations and standards of cassava cultural practices as outlined in previous studies [19,24,25].

#### 1.1 Phenotypic evaluation

Four architectural traits were measured including full plant height (PlantHeight), the first branching height (FirstBranchHeight), the branching level number (BranchlevelNum, Fig 1), and plant shape a categorical descriptor summarizing overall canopy architecture. The plant shape trait comprises four commonly recognized forms cylindrical, umbrella, compact, and open (Fig 2). Cylindrical types are characterized by the absence of branching, making them ideal for mechanized farming systems. Umbrella types typically branch once at a higher point (generally above 1 meter) and have relatively few subsequent branches. Compact and open types both branch at lower heights and develop multiple branching levels, but they differ in their architectural angles: compact plants exhibit more erect stems and narrower branch angles, while open types display a wider canopy spread due to broader branch divergence.

**Figure 1:**
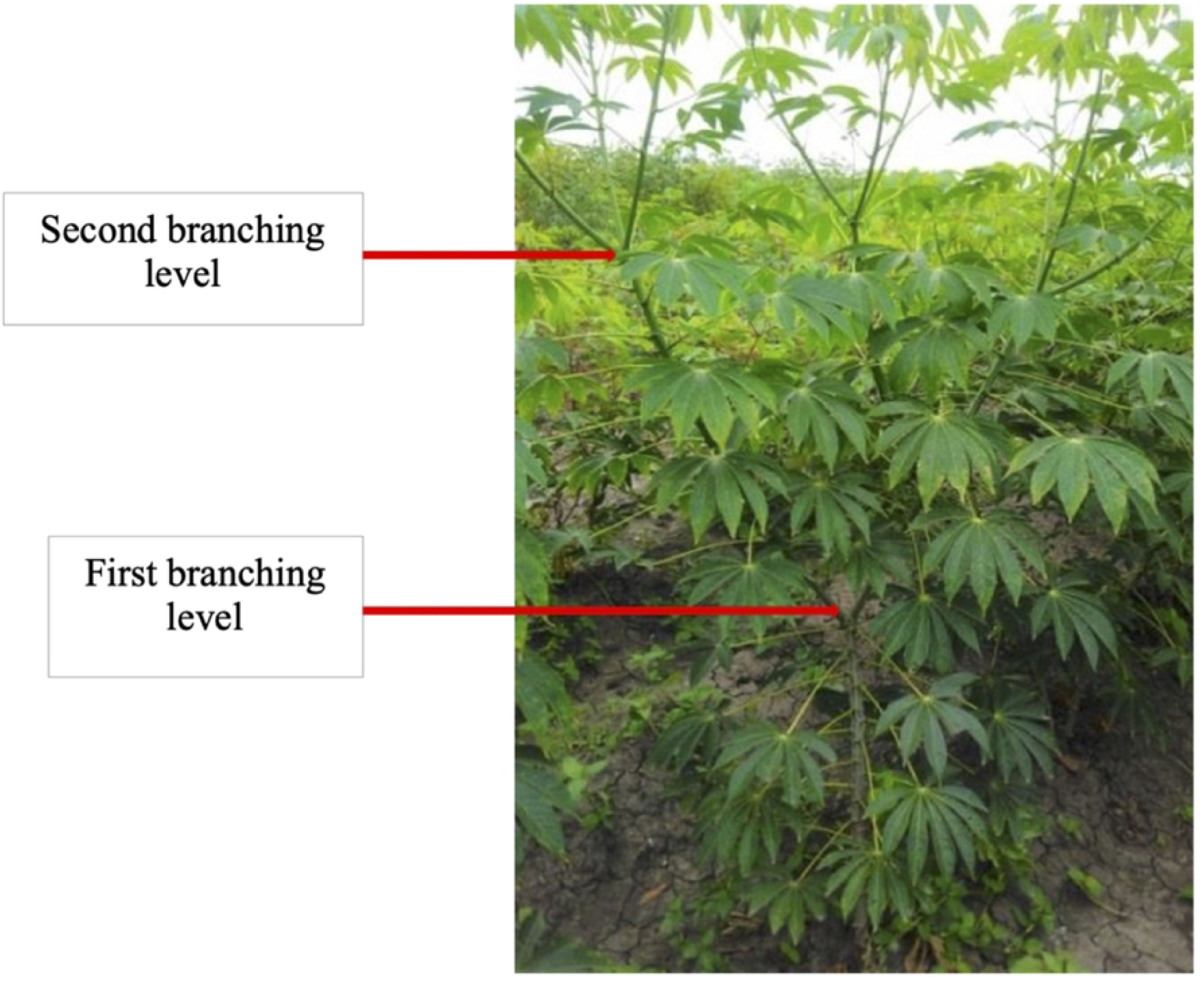
Cassava plant displaying two levels of branching (BranchlevelNum = 2). This figure illustrates a single plant with two distinct branching tiers along the main stem. In contrast, plants with BranchlevelNum = 1 typically exhibit a single primary branching point, while those with BranchlevelNum ***= 3*** develop a third tier of branching higher on the plant, resulting in a more complex canopy structure.

**Figure 2:**
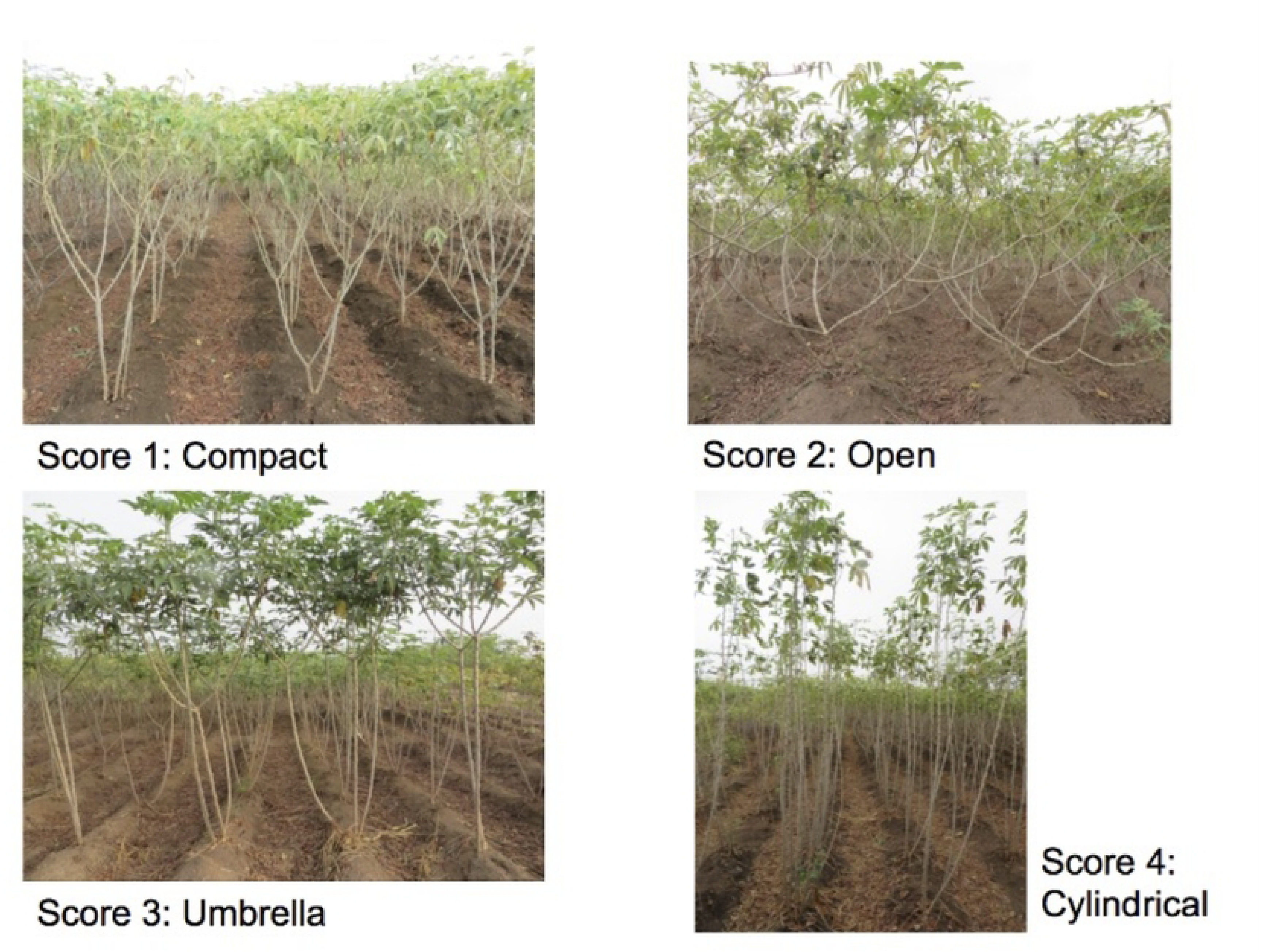
photos of the four most frequent cassava plant shape. Plant shapes differ primarily in FirstBranchHeight, BranchlevelNum, and canopy structure. Compact types show relatively short FirstBranchHeight, high branching density, and a dense canopy (Score 1). Open types exhibit very short FirstBranchHeight, high BranchlevelNum, and long, widely spaced branches forming a loose canopy (Score 3). Umbrella types are intermediate, with moderate FirstBranchHeight and branches concentrated in the upper canopy. Cylindrical types present tall FirstBranchHeight, minimal branching, and a narrow, erect canopy (Score 4).

All traits were observed at the plant level, but the final data were recorded as mean values per plot. Detailed trait descriptions and ontologies are available in (https://cassavabase.org/search/traits) and photos are provided in Fukuda et al. (2010). The data were collected at 9 months after planting (MAP).

The main traits of interest for which cassava is selected include fresh root yield (FYLD), above-ground biomass (TYLD) and dry matter content (DMC). In this study, correlation between these traits and those traits related to cassava plant architecture as mentioned above was investigated. The methods used to measure FYLD, TYLD and DMC were those used by Bakare et al. (2022).

### 2 Methods

#### 2.1 Single trials analysis

A preliminary analysis was performed for each trial to assess the relative strength of the genetic signal specifically heritability and reliability following the approach of Bakare et al. (2022) and Okoma et al. (2025). The analysis was conducted in two steps.

In the first step, the traits analyzed included PlantHeight, FirstBranchHeight, BranchlevelNum, FYLD, TYLD, and DMC. A linear mixed-effects model (lmer) from lme4 package was fitted in R 4.3.0 [28,29]. Given the diversity of experimental trial designs, two groups of trials were analyzed: replicated trials with at least two replicates and non-replicated trials with only incomplete block replicated controls.

- For replicated trials, the model formula was:

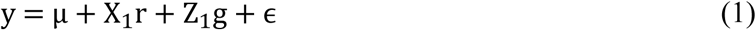

where y is the vector (n x 1) of observed agronomic trait, in which n is the number of observations in the trial; μ is the fixed intercept (overall mean); r is the (rx1) vector of fixed effects of replicates with its associated incidence matrix X_1_ of dimension nxr; g is the (gx1) vector of random effect of accession assumed to adhere to a Gaussian distribution 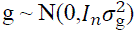 and its associated design matrix Z1 of dimension nxg and, ɛ represents a residual term assumed as 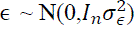.
- The Augmented Block Design analysis method (Kling, 2019) was used to fit the model for the non-replicated trials as follows:

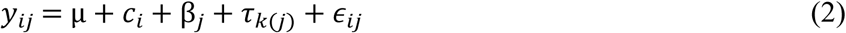

Three parameters were generated to perform this analysis:

- Entryc: Assigned a unique value to each check accession and a single, shared value for all test accessions. For instance, in a trial with 3 check accessions and 100 test accessions, the check values might be labeled “A”, “B”, and “C”, while all test accessions would be labeled “Z”;
- Entry: Assigned a unique identifier to each accession, including both checks and tests. Using the same example, identifiers would range from 1 to 103;
- New: A binary variable indicating accession type, with one level for checks and another for tests.

In Where:

- *y*_*ij*_ is a phenotypic vector of the observed agronomic trait of jth ‘check’ accession, in ith block;
- μ is the fixed intercept (overall mean);
- *c*_*i*_is a fixed effect assigned a unique value to each check accession and a single, shared value for all test accessions. For instance, in a trial with 3 check accessions and 100 test accessions, the check values might be labeled “A”, “B”, and “C”, while all test accessions would be labeled “Z”;
- β_*j*_ is the random effect of block j;
- τ_*k*(*j*)_is the random effect for “Entry:New” combination nested within block j where Entry is unique for each accession, including both checks and test accessions (In the example above, identifiers would range from 1 to 103), and New is a binary variable indicating accession type, with one level for checks and another for test accessions;
- ϵ_*ij*_ ϵ is the residual error term assumed to be normally distributed.

The second step of the analysis involved the plant shape variables including Cylindrical, Umbrella, Open, Compact. Because plant shape is categorical and the shapes cannot be ordered, each shape was analyzed as a separate binary variable (e.g., “cylindrical” versus “not-cylindrical”). In this framework, high heritability for a shape means that the shape is quite distinct (e.g., cylindrical accessions cannot be mistaken to have a different shape), whereas low heritability means that a shape can be confused with a different shape (e.g., open and compact might be interchangeable). Due to the binary nature of these variables, the generalized linear mixed-effects model (GLMM, lme4 package) was fitted by applying the binomial family function (link = “logit”) with the genotype effect set as random.

The genetic variance of each trial also was estimated using the procedure outlined by Bakare et al. (2022). The coefficient of variation (CV), broad-sense heritability (H^2^), and experimental accuracy (Ac) for all the traits were calculated for all traits in each trial using equations 3, 4, 5 respectively. To address potential outliers, the student function from the stats package was employed to identify and remove row data with Studentized Residuals exceeding 3 [29]. A total of 257 trials across 28 locations, involving 9,062 accessions, were retained for further analysis based on criteria of ≥ 50% experimental accuracy (Ac) and broad-sense heritability (H²) > 0.1 for at least one trait (Table 1).

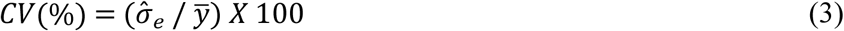

**Table 1:** Summary of plant architecture traits, experimental designs, and germplasm evaluated across Nigerian cassava breeding programs.

| Trait | Breeding program | Trial design | Year | Location | #Trials | #accessions | Ontology |
| --- | --- | --- | --- | --- | --- | --- | --- |
| PlantHeight | IITA | Alpha | 4 | 12 | 37 | 2080 | Plant height measurement (cm) |
| PlantHeight | IITA | Augmented | 3 | 2 | 3 | 758 |  |
| PlantHeight | IITA | CRD | 3 | 4 | 5 | 109 |  |
| PlantHeight | IITA | RCBD | 9 | 22 | 87 | 1813 |  |
| PlantHeight | NRCRI | Alpha | 5 | 3 | 12 | 964 |  |
| PlantHeight | NRCRI | Augmented | 2 | 2 | 3 | 581 |  |
| PlantHeight | NRCRI | RCBD | 6 | 7 | 50 | 679 |  |
| FirstBranchHeight | IITA | Alpha | 4 | 10 | 34 | 2042 | First apical branch height measurement (cm) |
| FirstBranchHeight | IITA | Augmented | 4 | 3 | 4 | 835 |  |
| FirstBranchHeight | IITA | CRD | 2 | 2 | 2 | 49 |  |
| FirstBranchHeight | IITA | RCBD | 9 | 12 | 66 | 1909 |  |
| FirstBranchHeight | NRCRI | Alpha | 4 | 3 | 7 | 594 |  |
| FirstBranchHeight | NRCRI | Augmented | 1 | 1 | 3 | 419 |  |
| FirstBranchHeight | NRCRI | RCBD | 5 | 5 | 30 | 266 |  |
| BranchlevelNum | IITA | Alpha | 4 | 9 | 31 | 2289 | Branching level counting |
| BranchlevelNum | IITA | Augmented | 4 | 3 | 5 | 1348 |  |
| BranchlevelNum | IITA | CRD | 2 | 2 | 2 | 49 |  |
| BranchlevelNum | IITA | RCBD | 8 | 7 | 39 | 1357 |  |
| BranchlevelNum | NRCRI | Alpha | 1 | 1 | 1 | 25 |  |
| PlantShape | IITA | Alpha | 2 | 1 | 5 | 163 | Plant shape visual assessment |
| PlantShape | IITA | Augmented | 2 | 3 | 4 | 1764 |  |
| PlantShape | IITA | RCBD | 2 | 4 | 9 | 140 |  |

This table includes 9,062 accessions retained after single-trial filtering for phenotypic analysis. Columns indicate: Trait (plant architecture trait evaluated); Breeding program (IITA or NRCRI); Trial design (experimental setup); Year (number of distinct years of evaluation); Location (trial site); #Trials (number of trials per trait); #Accessions (unique genotypes evaluated); and Ontology (standardized trait definition and measurement).

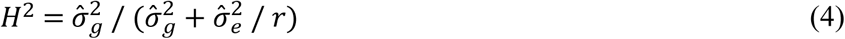

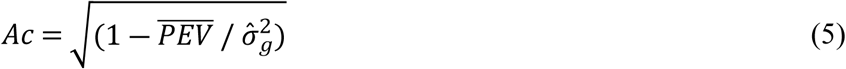

where 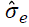 is the model residual standard deviation, 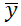 is the estimate of the overall mean for a given trait, 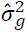 is the estimated genetic variance (clonal variance), 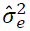 is the model residual variance, *r* is the number of replicates and, 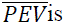 the average of prediction error variance.

#### 2.2 Multiple trials analysis

The distribution of each variable among the breeding program including the breeding stage was defined. The best linear unbiased predictor (BLUP) was obtained for each trait and accession by fitting the models cited above respectively to the type of trait. For the first group of variables, a single lmer model was fitted as presented in Eq2, including both repeated and unrepeated trials. Prediction error variance (PEV) of each accession was extracted and the deregressed BLUP (drgBLUP) was estimated according to Eq6, following the method of Garrick et al. (2009). Trait correlations were then computed using the resulting drgBLUP.

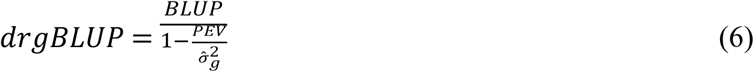

○ PEV is the prediction error variance of each accession
○ 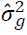 accessions variance

#### 2.3 Genotyping, SNPs quality control, Population Structure, and Linkage Disequilibrium Analysis

DNA extraction, genotyping-by-sequencing (GBS), single nucleotide polymorphism (SNP) calling, and marker quality control were performed following the procedures described by Wolfe et al. (2017); Rabbi et al., (2014, 2017) and Olayinka et al., (2025). A total of 4,530 accessions were phenotyped and genotyped, yielding 61,239 high-quality SNPs with a minor allele frequency (MAF) > 0.01 were used for downstream analyses. Population structure was evaluated prior to genome-wide association analysis to control for confounding due to genetic stratification within the panel. Principal Component Analysis (PCA) was performed on the SNP dosage matrix using the prcomp function in R [29].

Genome-wide linkage disequilibrium (LD) was characterized to describe the extent of marker correlation and to inform downstream association analyses. Pairwise LD was estimated as the intrachromosomal squared correlations coefficient (*r²*) among 61,239 SNPs using PLINK v1.9 [35], considering all marker pairs within a 1 Mb physical distance window. LD decay was visualized in R [29] by plotting r² values against inter-marker physical distance on a log₁₀ scale, using hexagonal binning to summarize SNP-pair density and a smoothed trend line to represent the mean LD decay pattern across distances.

#### 2.4 GWAS Analyses

Genome-wide association studies (GWAS) were performed using the Genomic Association and Prediction Integrated Tool (GAPIT 3) package in R [29]. To ensure comparability with previous cassava studies while maximizing detection power, two complementary GWAS models were applied. Model selection was informed by the genome-wide LD profile of the panel, and both a mixed linear model (MLM) and the BLINK algorithm were implemented to accommodate the observed LD structure and mitigate confounding. The MLM model was implemented as the reference approach because it has been widely adopted in cassava GWAS to control for population structure and relatedness through a unified kinship (K) matrix and principal components (PCs). In addition, we used the Bayesian-information and Linkage-disequilibrium Iteratively Nested Keyway (BLINK) model, which extends the multi-locus framework to account for linkage disequilibrium and iteratively refine associated markers, thereby increasing statistical power without inflating false positives.

Analyses were performed using drgBLUPs as the response variable for all the traits. For both models, markers with minor allele frequency (MAF) < 0.01 were excluded, and the first three PCs were included to correct for residual population structure. Genome-wide and suggestive significance thresholds were set at −log₁₀(0.05/M) and −log₁₀(1/M), respectively, where M = 61,239 SNPs. Manhattan and quantile–quantile (Q–Q) plots were generated to visualize the SNPs–trait associations and to assess the overall fit and inflation of the test statistics.

#### 2.5 Genomic Prediction

Genomic best linear unbiased prediction (G-BLUP) was performed to estimate Genomic Estimated Breeding Values (GEBV) for the studied population. This was done using the sommer R package [36].

The study of Andrade et al. (2022) showed that for a given trait, G-BLUP accuracy can vary according to the type of genetic models: additive or additive-dominance. Based on these results, these two models were evaluated for their cross-validation accuracy. The G-BLUP model fitted was as following:

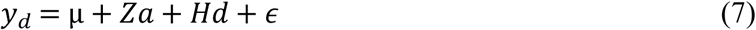

where *Yd* is the vector of the drgBLUP, μ is the overall mean, a vector of additive effect 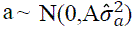, d is the dominant deviation effect with 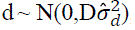, Z and H are the incidence matrices for μ, a and d, respectively, with 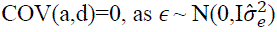. The additive relationship matrix A and the classical dominant relationship matrix D were constructed using the package genomicMateSelectR in R [29]

K-fold cross-validation was used to assess the accuracy of predicting the genomic estimated breeding values (GEBVs) for candidate parents lacking phenotypic data. The analysis was performed using 5 folds with three repetitions. Prediction accuracy for each fold was calculated as the Pearson correlation between the GEBVs of the validation set and the corresponding BLUPs derived from the test sets.

## Results

### 1 Phenotype analysis: Statistical Summary, Variability, Correlations, and Heritability

The phenotypic data revealed a highly significant difference in how ongoing selection has influenced plant architecture between IITA and NRCRI. As illustrated in Fig 3, PlantHeight at IITA showed a significant reduction (p < 0.001), decreasing from an average of 1.96 m in CET to 1.63 m in UYT. In contrast, the opposite trend was observed at NRCRI, where the average height increased from 1.66 m in CET to 1.87 m in UYT. A similar pattern was noted for FirstBranchHeight in each breeding program. The results of the ANOVA test are provided in the supplementary Table S1. The average BranchlevelNum measured mainly in IITA increased significantly from 2 at CET to 3 at UYT. The distribution of residuals reflects the normal distribution of the first group of traits (S1 Fig).

**Figure 3:**
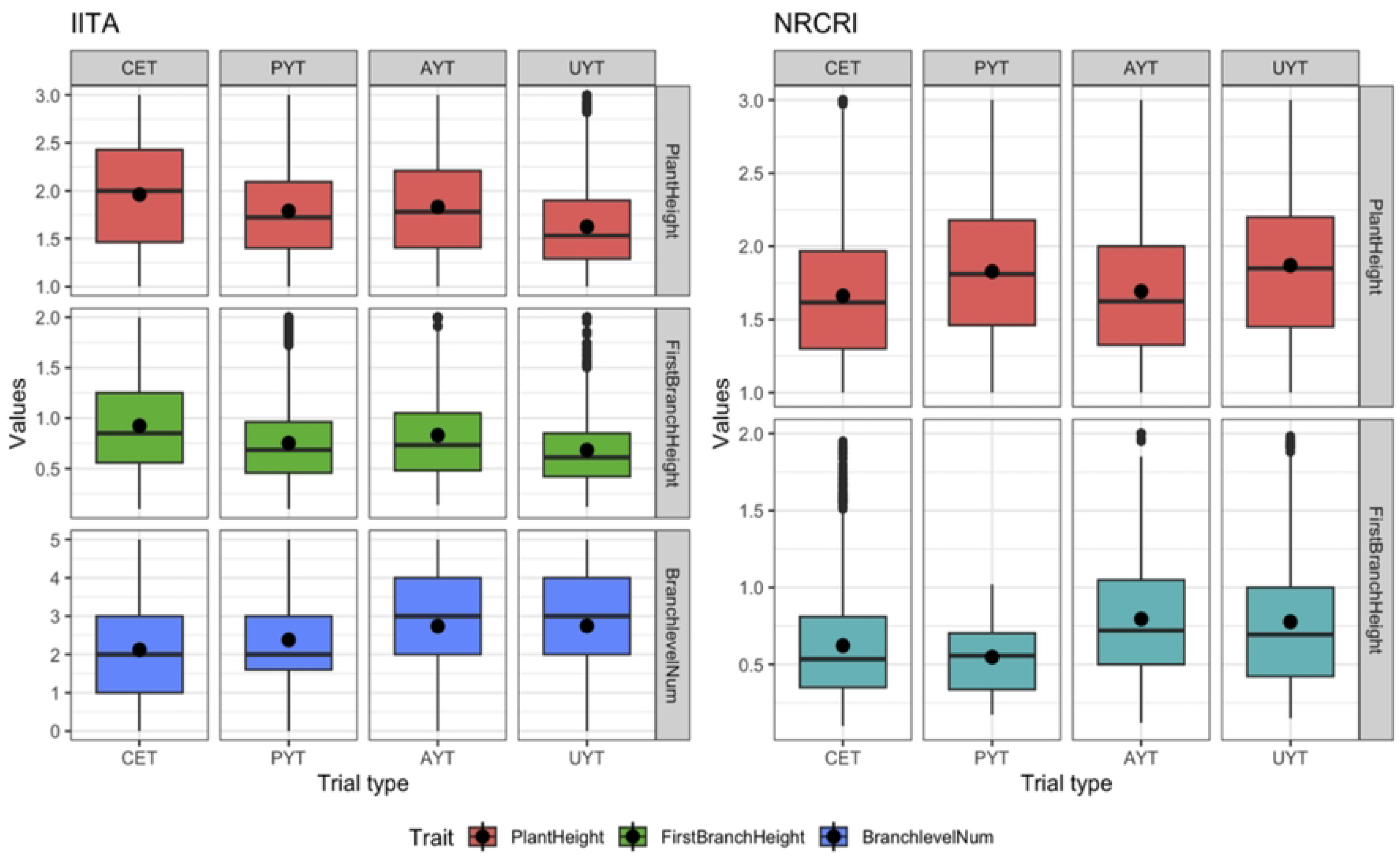
Distribution of PlantHeight, FirstBranchHeight and BranchlevelNum across breeding stages in Nigeria Cassava germplasm from IITA and NRCRI.

The distribution of plant shape traits indicates that the IITA breeding population was predominantly characterized by the open type from the CET to the AYT stages, accounting for 42, 46, and 52%, respectively, for CET, PYT, and AYT (Fig 4). Subsequently, the population of the open type declined to 7% at the UYT stage, giving way to increased frequencies of the compact and cylindrical types. The last stage (UYT) was marked by the predominance of the umbrella type, comprising 44% of the population.

**Figure 4:**
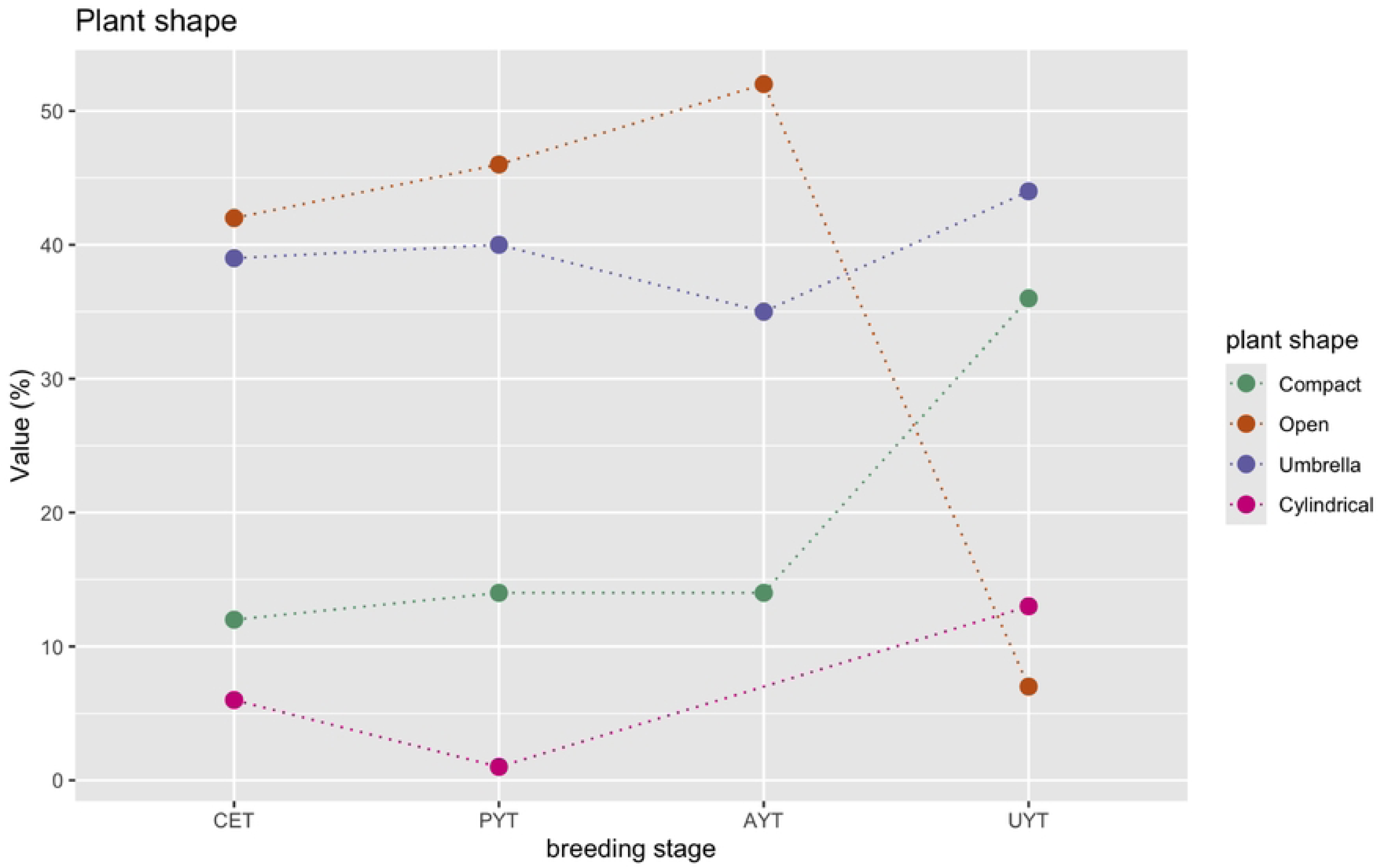
Distribution of Compact, Open, Umbrella and Cylindrical type throughout the breeding stages of IITA germplasm.

Broad-sense heritability (H^2^) for all the traits ranged from 0.41 to 0.72 (Table 2). Notably, BranchlevelNum exhibited the highest H^2^ within the first group of traits, reaching 0.65. Furthermore, the values of H^2^ for PlantHeight and FirstBranchHeight were closely comparable and moderate, 0.52 and 0.55, respectively. For the plant shape traits, Cylindrical demonstrated the highest H^2^ at 0.72, while Open showed the lowest H^2^ at 0.41. Additionally, the heritability values for Compact and Umbrella, 0.66 and 0.55 respectively, are noteworthy and contribute to the overall understanding of trait heritability in this study.

**Table 2:**
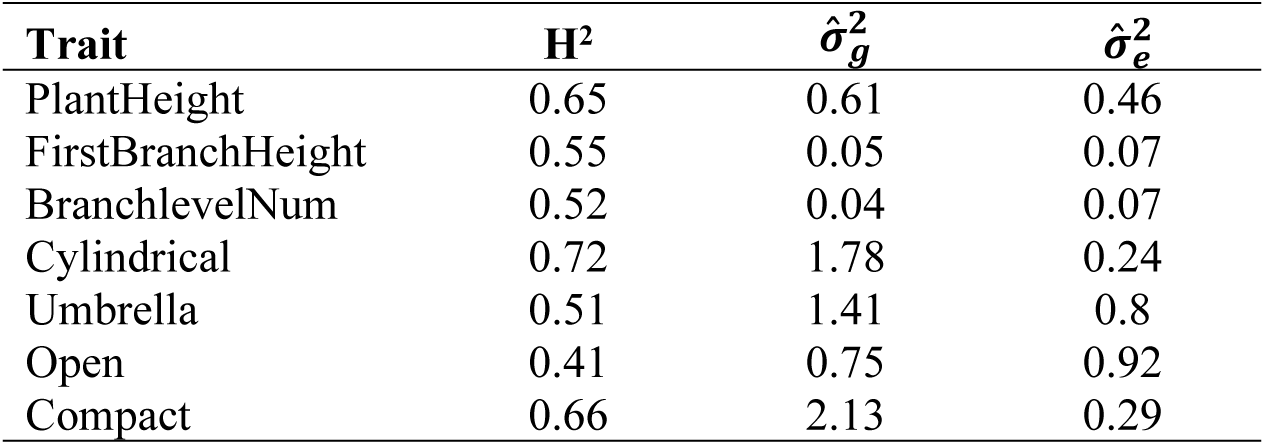
Broad sense heritability and Variance components for Cassava architecture Traits.

The correlation between plant architectural traits and general yield trait such as dry matter content (DMC), fresh root yield (FYLD), and above-ground biomass (TYLD) was investigated (Fig 5). Many of the architectural traits exhibited negligible correlation, with respect to the yield traits, except for PlantHeight, which showed a slight positive correlation with TYLD (0.32). The drgBLUP for BranchlevelNum exhibited a slight yet statistically significant negative correlation with both FirstBranchHeight (−0.40) and cylindrical types (−0.35). A slight positive correlation was observed between BranchlevelNum and the open type (0.29), the latter displaying a notable negative correlation with the umbrella type (−0.64). On other hand, PlantHeight had slight positive correlation with FirstBranchHeight (0.36).

**Figure 5:**
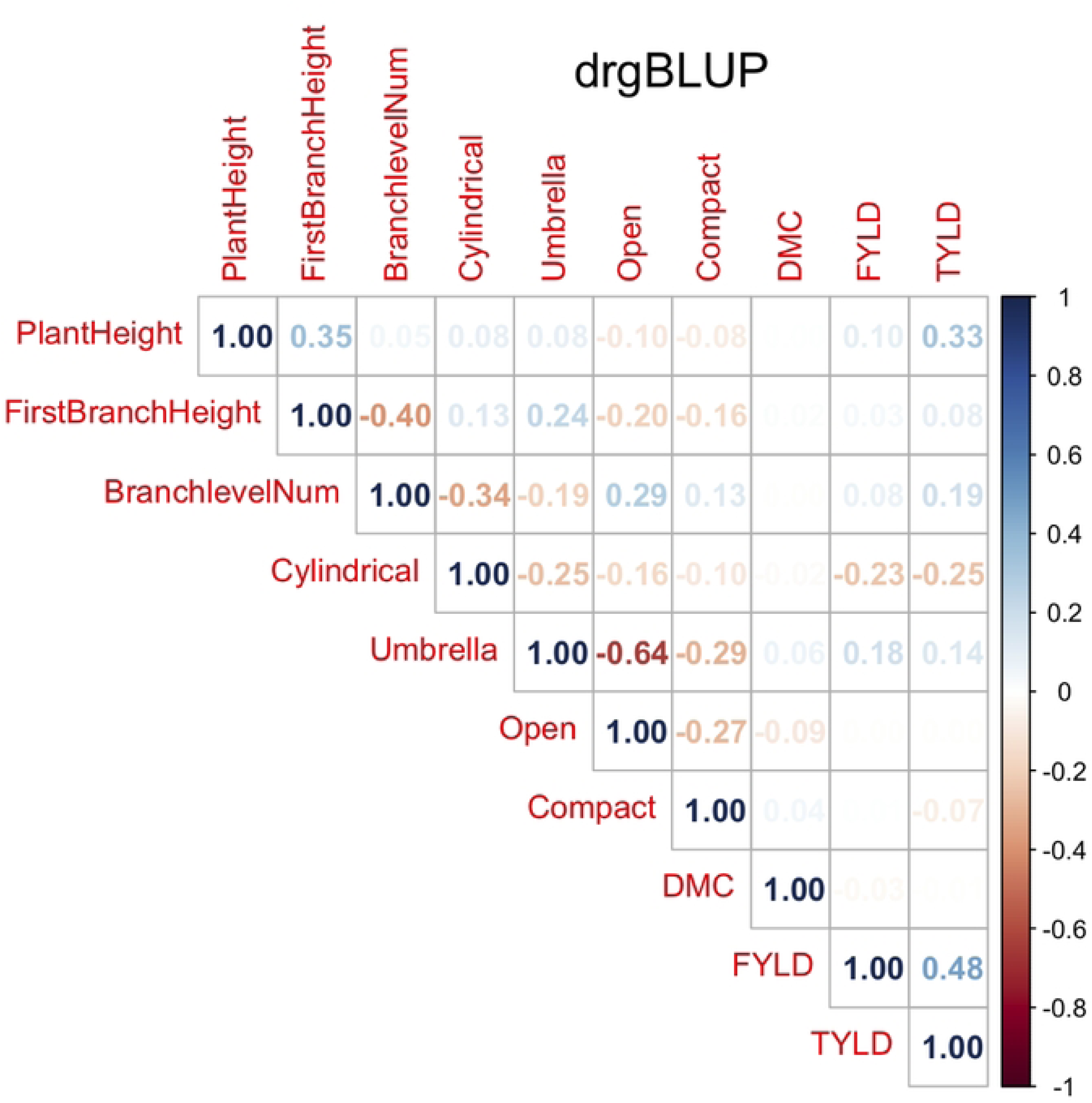
Heatmap of pairwise trait correlations using drgBLUP from 10 traits including 7 architecture traits and 3 yield component traits dry matter content (DMT), fresh storage root weight (FRWT) and fresh shoot weight (FTWT)

### 2 Genomic analysis and GWAS

#### 2.1 Population Genetic structure and Linkage Disequilibrium Patterns

The first two principal components PC1 and PC2, which accounted for 10 % of the total genetic variations (Fig 6 and S2 Fig) were used to visualize population structure. Principal component analyses on the SNP marker matrix did not detect subtle genetic differentiation among the NRCRI and IITA (Fig 6). Further, the plot of the contribution values of each marker on PC1 and PC2 against marker positions along the 18 cassava chromosomes revealed that the markers with the strongest influence on PC1 were located on the first chromosomes (Fig 7).

**Figure 6:**
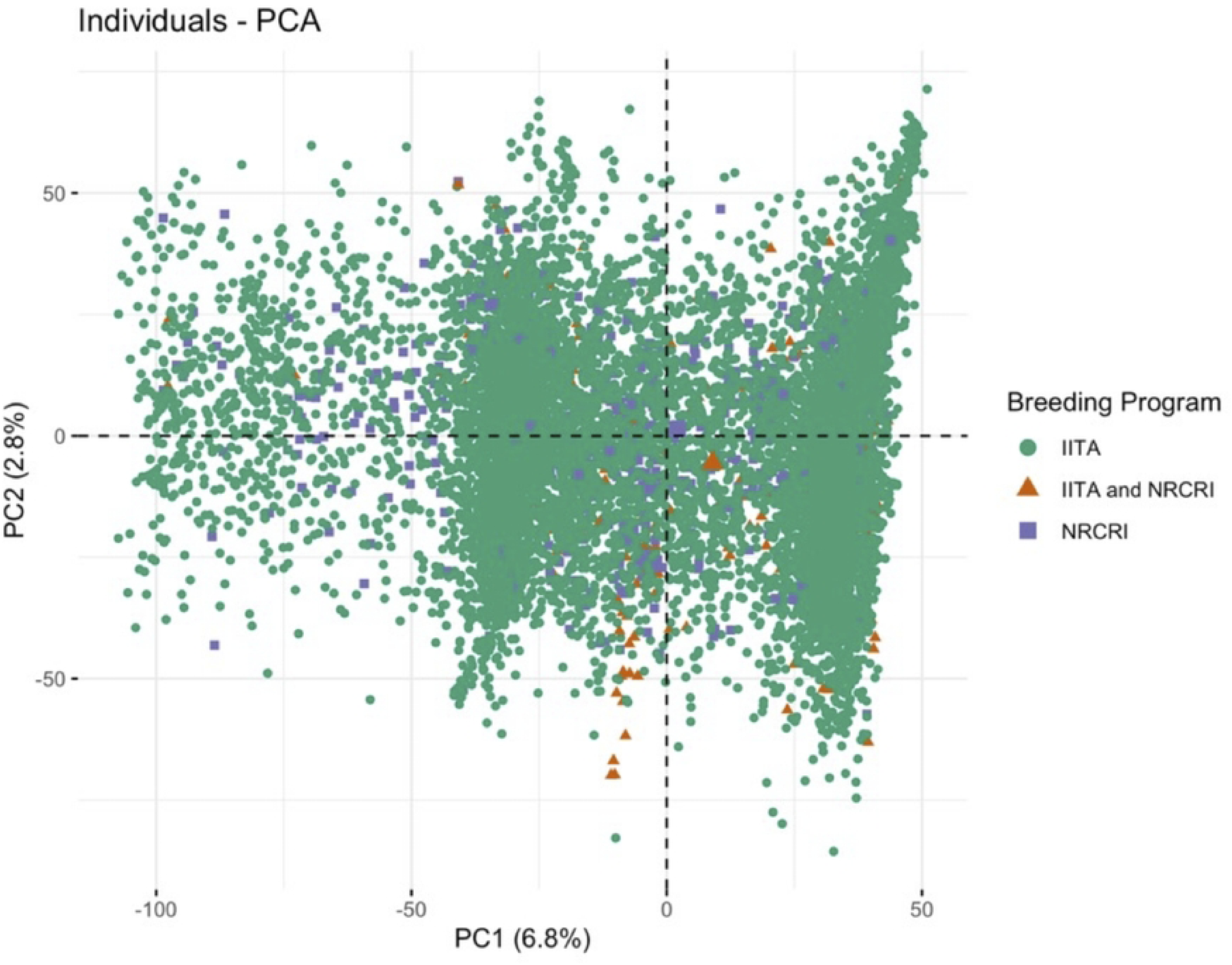
Population structure of the Genetic Gain collection as revealed by the first two axes of principal component analysis (PCA) biplot. Green circles indicate individuals from the IITA program, Orange triangles represent individuals shared between IITA and NRCRI and Purple squares represent individuals exclusive to NRCRI

**Figure 7:**
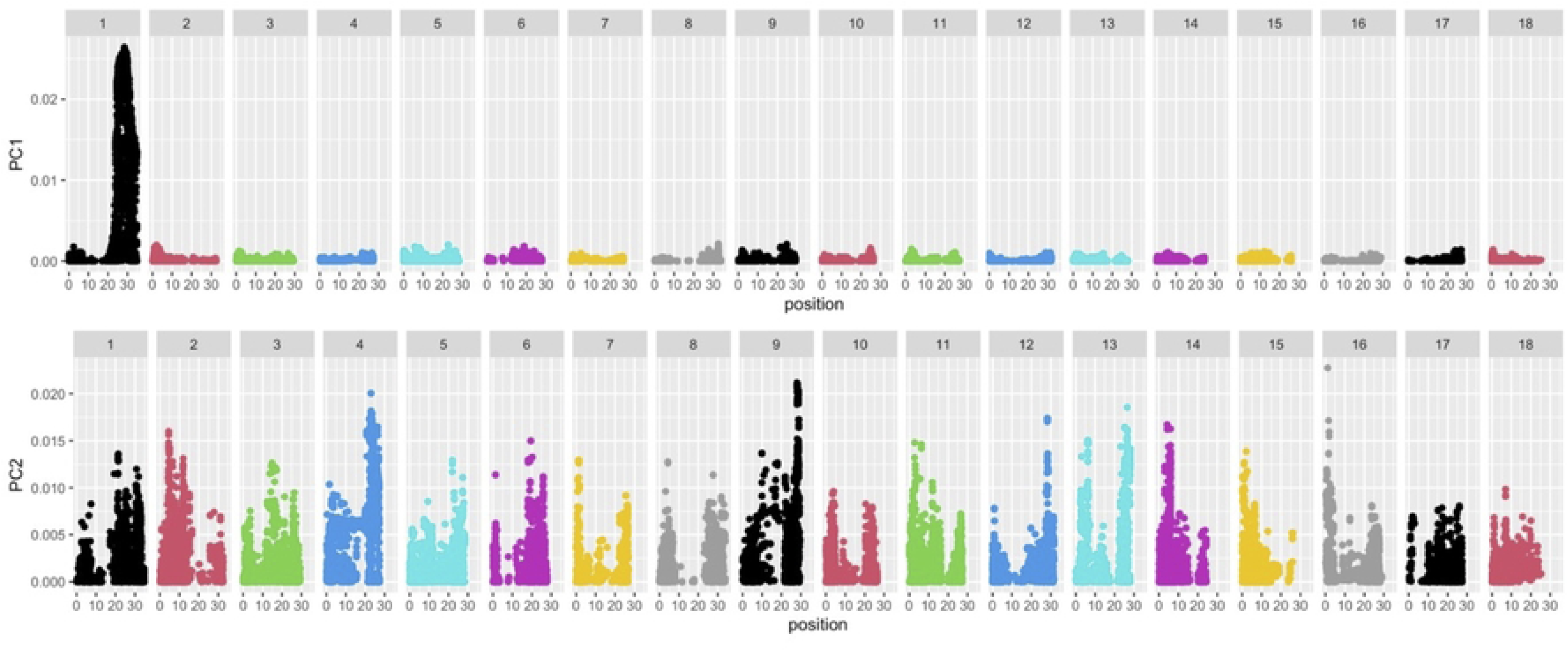
Plot of each marker’s contribution to Principal Component 1 (PC1) and Principal Component 2 (PC2) against their positions along the 18 cassava chromosomes revealed that the markers most strongly influencing PC1 were located on the first chromosome.

Genome-wide linkage disequilibrium (LD) analysis revealed a characteristic decay pattern across the cassava genome (Fig 8). Pairwise LD (r²) was highest at short inter-marker distances, reaching values around 0.44 for closely linked SNPs, followed by a rapid initial decline within the first few kilobases. Beyond this early drop, LD decreased more gradually with increasing physical distance, approaching r² values near 0.20 at distances close to 1 Mb.

**Figure 8:**
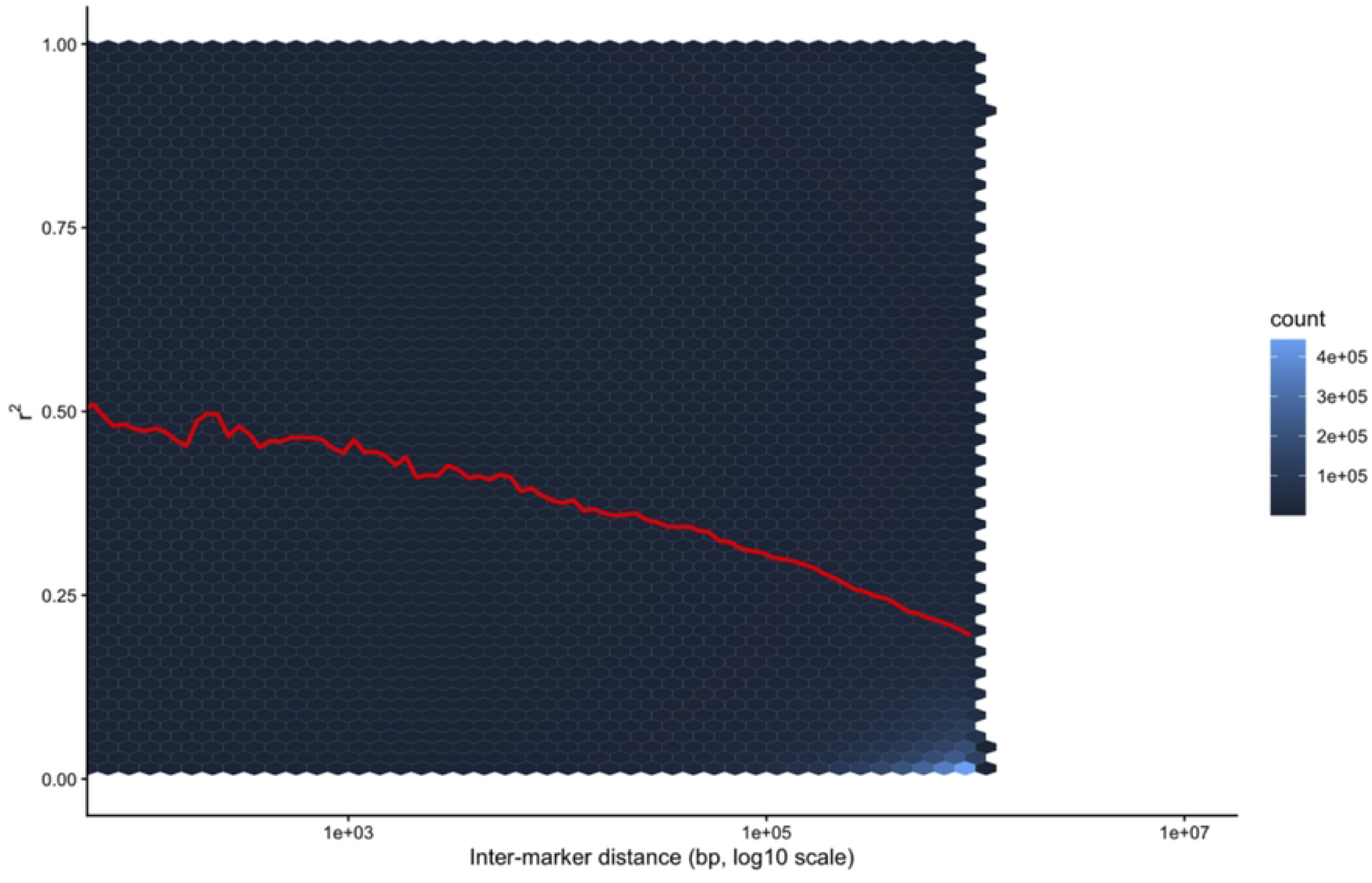
Genome-wide linkage disequilibrium (LD) decay in cassava. Pairwise LD estimates (r²) were calculated for all intrachromosomal SNP pairs within a 1 Mb physical distance window. Hexagonal bins represent the density of SNP pairs as a function of inter-marker distance plotted on a log₁₀ scale. The black line shows the smoothed mean LD trend across distance. LD declines sharply at short distances and decreases gradually over longer genomic intervals, reflecting extended haplotype structure in the cassava breeding panel.

#### 2.2 GWAS detection of a shared QTLs for BranchlevelNum

Genome-wide association analysis involving 3,113 accessions with complete phenotypic records for all architectural traits and 61,239 SNPs revealed both shared and model-specific association patterns across the cassava genome. Although GWAS was performed for all evaluated architectural traits, BranchlevelNum was the only trait for which both BLINK and MLM detected a concordant genomic region exceeding the significance threshold. BLINK identified multiple significant loci across the genome, with its strongest association observed on chromosome 2 at SNP *S2_2809137*, which accounted for 5.8% of the phenotypic variance. In contrast, MLM yielded a more conservative set of associations confined to a compact cluster of eight SNPs between 2.80 Mbp and 3.00 Mbp on chromosome 2, among which *S2_2846222* and *S2_2854244* explained approximately 14.44% and 10.0% of the phenotypic variance, respectively (Table 3). These SNPs showed consistent effect sizes and directions across models and formed a single, sharply defined association peak, indicating that they represent multiple markers in strong linkage disequilibrium tagging the same underlying QTL for BranchlevelNum. Consistent with this interpretation, pairwise LD estimates among the significant SNPs within the chromosome 2 QTL showed uniformly high r² values (median r² ≈ 0.77), supporting the presence of a single underlying locus (S2 Table). Despite differences in model sensitivity, both BLINK and MLM converged on the same chromosome 2 interval as the principal locus underlying variation in BranchlevelNum.

**Table 3:** Summary of significant SNPs associated with Number of Branching Levels identified by GWAS using a Mixed Linear Model.

| SNP | Chromosome | Position | P.value | maf | nobs | Rsquare.of.<br>Model.<br>without.SNP | Rsquare.<br>of.Model.<br>with.SNP | effect | Phenotype<br>_Variance<br>Explained(%) |
| --- | --- | --- | --- | --- | --- | --- | --- | --- | --- |
| S2_2809137 | 2 | 2809137 | 2.91E-07 | 0.06087376 | 3113 | 0.17006362 | 0.17714039 | -0.193331 | 1.01E-07 |
| S2_2818574 | 2 | 2818574 | 2.91E-07 | 0.06087376 | 3113 | 0.17006362 | 0.17714039 | -0.193331 | 8.62E-08 |
| S2_2828773 | 2 | 2828773 | 2.74E-07 | 0.06103437 | 3113 | 0.17006362 | 0.17717218 | -0.1937591 | 0 |
| S2_2846222 | 2 | 2846222 | 6.28E-07 | 0.06023129 | 3113 | 0.17006362 | 0.17674 | -0.1907898 | 2.571682981 |
| S2_2854244 | 2 | 2854244 | 4.82E-07 | 0.06151622 | 3113 | 0.17006362 | 0.17687752 | -0.1886327 | 4.45E-08 |
| S2_3000720 | 2 | 3000720 | 1.68E-07 | 0.06537102 | 3113 | 0.17006362 | 0.17742618 | -0.1900843 | 2.192235115 |
| S13_266164 | 13 | 266164 | 5.39E-07 | 0.23931898 | 3113 | 0.17006362 | 0.17681956 | -0.1096813 | 6.028071398 |

Detailed information on significant SNPs and their genomic positions is provided in Table 3. Manhattan and quantile–quantile (Q–Q) plots for BranchlevelNum, illustrating the distribution of observed versus expected association statistics, are shown in Fig 9, and the corresponding plots for the remaining traits are presented in S3 Fig.

**Figure 7:**
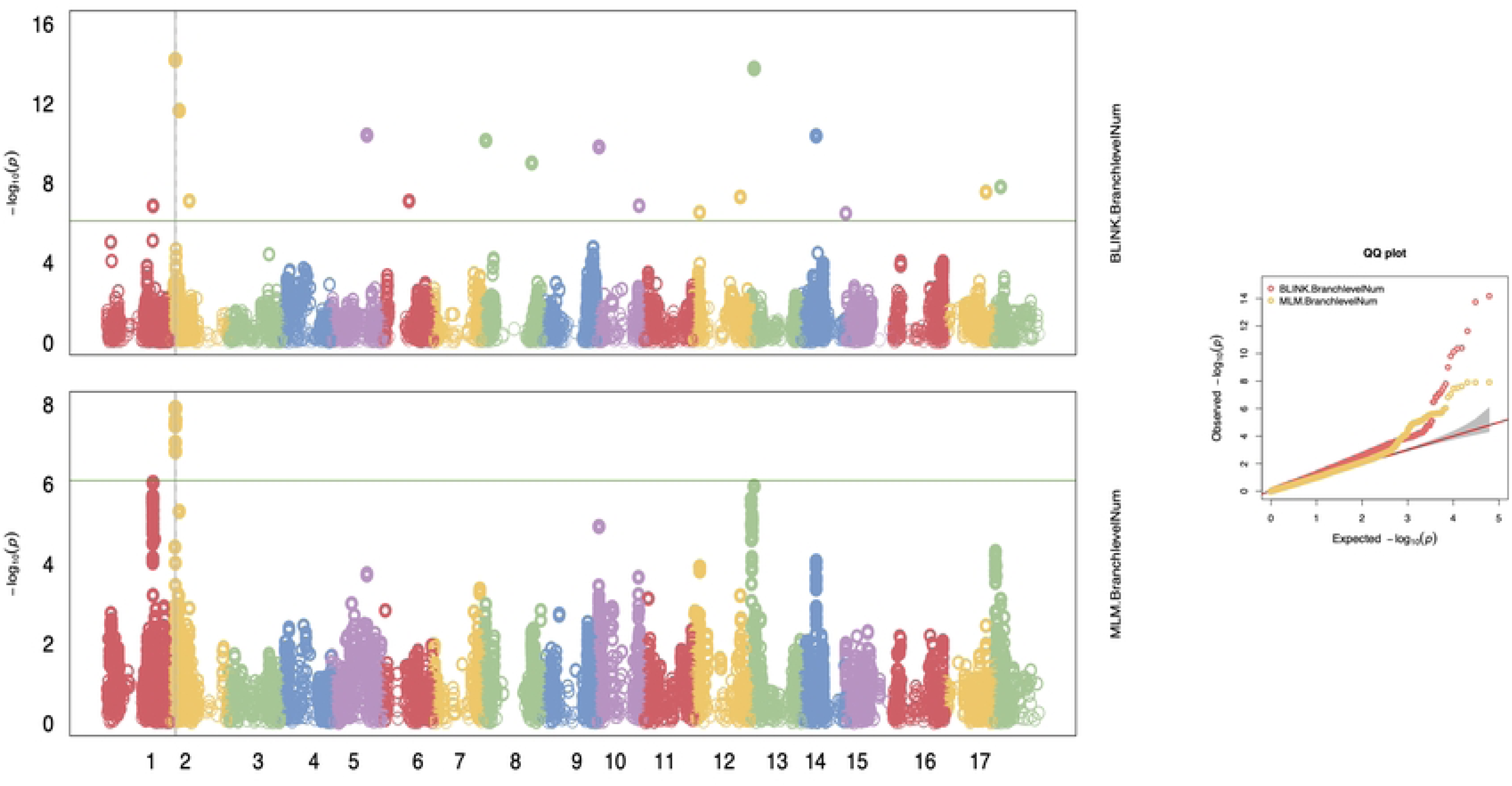
Manhattan plots (left) and corresponding quantile-quantile (Q-Q) plots (right) of the Bayesian-information and Linkage-disequilibrium Iteratively Nested Keyway (BLINK) model and the mixed linear model (MLM) results for plant height architecture traits across 18 chromosomes in 3,113 cassava accessions from the IITA and NRCRI breeding programs. In the Manhattan plot, the X axis is the genomic position of the SNPs in the genome, and the Y axis is the negative log base 10 of the P-values. Each chromosome is colored differently. SNPs with stronger associations with the trait will have a larger Y-coordinate value. the solid green horizontal line indicates the significance threshold at −log₁₀(P).

#### 2.3 Identification of Putative Candidate Genes for BranchlevelNum

Putative candidate genes underlying variation in BranchlevelNum were identified by defining a candidate genomic interval on chromosome 2 based on the physical positions of the three SNPs explaining more than 5% of the phenotypic variance. This candidate genomic region spanned approximately 2.80–2.90 Mbp on chromosome 2. Genes located within this region were retrieved using functional annotations from Phytozome v13 (Manihot esculenta v7.1, accessed December 28, (*2024*) and the National Center for Biotechnology Information (NCBI) cassava genome database (accessed December 28, (*2024*).

A total of 16 annotated genes associated with cassava branching phenology were identified (S2 Table). Genes with known functions in plant development and phenology were grouped into three functional categories. The first category comprised genes involved in organ growth, development, and cell division, including *Manes.02G025755*, homologous to SCARECROW-LIKE 28, and *Manes.02G027800*, which encodes a MATE efflux family protein implicated in developmental regulation [40]. A second category included genes associated with reproductive organ development and fertility, such as *Manes.02G025740*, homologous to Protein Hapless 2, which is required for pollen tube guidance and successful fertilization[41]. The third category consisted of genes encoding cytochrome P450 enzymes, which are widely involved in hormone metabolism, developmental processes, and defense responses in plants; notable examples include *Manes.02G025745* and *Manes.02G025760* [42,43].

### 3 Genomic Prediction Accuracy for Plant Architecture Traits

For each trait, the ANOVA mean comparison test showed no significant difference between the two genomic prediction models: additive and additive-dominance (Table 4).

**Table 4:** ANOVA Model and Summary of Results Comparing Genomic Prediction Accuracy Across Seven traits.

| **Model: aov(Accuracy ~ Model\*Trait)** |  |  |  |  |  |
| --- | --- | --- | --- | --- | --- |
|  | Df | Sum.Sq | Mean.Sq | F.value | Pr(>F) |
| Model | 1 | 0.000 | 0.0001 | 0.064 | 0.801 |
| Trait | 6 | 5.474 | 0.9124 | 438.074 | <2e-16 *** |
| Model:Trait | 6 | 0.001 | 0.0002 | 0.083 | 0.998 |
| Residuals | 297 | 0.619 | 0.0021 |  |  |

BranchlevelNum was the best trait for predicting plant architecture based on the genome, with an accuracy of 0.44 (Fig 10). PlantHeight and FirstBranchHeight had similar prediction accuracies of 0.25 each. The results also indicated that the Open_type trait had a comparable predictive potential to PlantHeight and FirstBranchHeight, with an accuracy of 0.26. Genomic prediction accuracy for other architectural parameters varied, ranging from 0.02 for Umbrella_type to 0.16 for Cylindrical_type.

**Figure 8:**
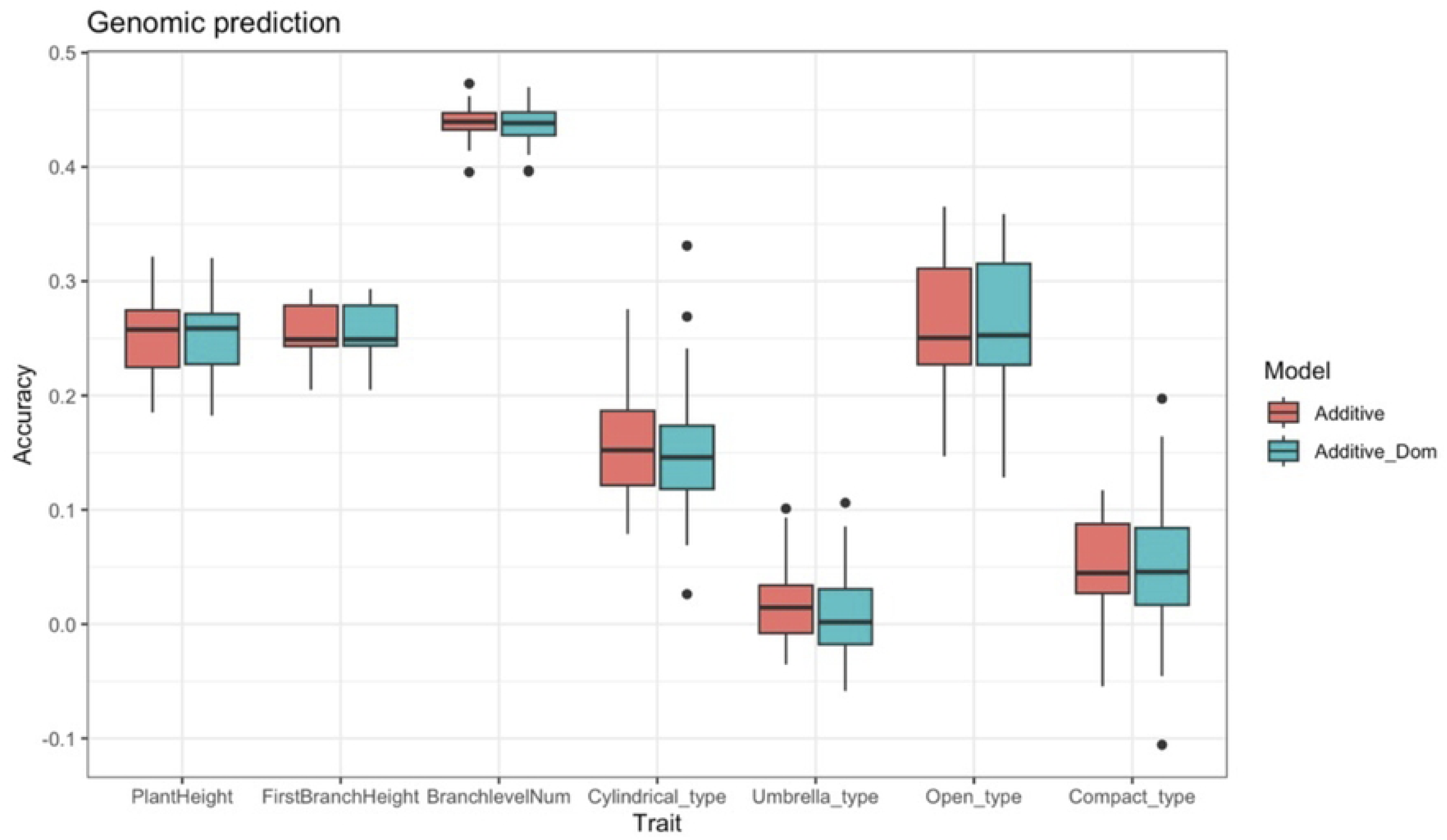
Comparison of Genomic Prediction Accuracy across seven traits using Additive and Additive-Dominance Models.

## Discussion

### 1 Selection for plant architecture in Nigerian Cassava breeding programs: a comparative perspective

The phenotypic analysis across selection stages indicated that plant architecture is selected upon differently within the two Nigerian breeding pipelines. At NRCRI, PlantHeight and FirstBranchHeight increased across successive breeding stages, whereas in the IITA program both traits decreased and BranchlevelNum increased. Because these patterns were derived directly from multi-year, multi-location phenotypic summaries, they provide evidence that the two programs progress with distinct architectural outcomes, even though they operate within broadly similar agro-ecological contexts.

We did not identify a consistent genetic correlation between yield and architectural traits across environments, cautioning against attributing architectural change directly to yield selection. Previous studies nevertheless provide important context for interpreting these trends. Olayinka et al. (2025) reported a negative and significant correlation between plant height and harvest index, with the strength of this relationship varying across environments, while Uwah et al. [44] demonstrated that high nitrogen application can increase both plant height and tuber yield. Together, these findings underscore the potential for environmental and agronomic factors to influence plant architecture and reduce genetic associations between morphology and productivity. As a result, interpretations of architectural change as a direct response to yield selection should be regarded as hypothesis-generating, highlighting the need to distinguish environmentally mediated effects from genetically regulated variation. Despite their contrasting early-stage trajectories, both breeding programs ultimately converged on similar plant architecture types at the release stage.

Within the IITA pipeline, this convergence was reflected by a shift in plant shape composition, with “open” architectures predominating in early-generation trials (CET to AYT) and umbrella and compact architectures becoming more frequent in later stages, particularly the Uniform Yield Trials (UYT). Similarly, several varieties released by the NRCRI program such as Game-Changer, Hope, and Obasanjo-2 have been described as exhibiting profuse or mixed branching patterns consistent with compact or umbrella architectures [45]. This convergence suggests that different combinations of PlantHeight, FirstBranchHeight, and branching intensity can lead to functionally equivalent canopy forms that satisfy late-stage selection criteria.

### 2 Heritability and trait relationships: implications for selection

Moderate to high broad-sense heritability estimates (H² = 0.41–0.72) observed indicate that several cassava architectural traits are under substantial genetic control and are therefore amenable to selection. In particular, Cylindrical type (H² = 0.72) and BranchlevelNum (H² = 0.65) exhibited high heritability, highlighting their potential as stable targets for early-generation selection and genomic prediction. These estimates are comparable to those reported by Olayinka et al., (2025), who observed heritability values ranging from 0.70 to 0.90 in more spatially and temporally restricted trials, suggesting that the genetic determinism of these traits is robust across diverse environments and experimental designs.

Among the architectural traits evaluated, BranchlevelNum stands out for both its breeding utility and biological relevance. Its established association with flowering capacity supports its role in improving reproductive synchrony and crossing efficiency [9,12,20]. Reported heritability estimates for BranchlevelNum vary from moderate to highly stable across environments consistent with the relatively stable inheritance observed.

PlantHeight and FirstBranchHeight showed moderate heritability (H² = 0.52 and 0.55, respectively), indicating that although these traits are genetically tractable, they are more strongly influenced by environmental conditions. This aligns with previous findings that architectural traits often exhibit substantial phenotypic plasticity due to their polygenic nature and sensitivity to management and environmental variation [14,23,46]. These patterns support a tiered breeding strategy, in which highly heritable traits such as BranchlevelNum and Cylindrical type are prioritized for early selection or genomic prediction, while more environmentally responsive traits are refined through multi-environment testing or later selection stages.

Trait correlation analyses further inform breeding decisions. Most architectural traits showed weak or negligible correlations with yield components such as fresh root weight and dry matter content, suggesting relative genetic independence and enabling simultaneous optimization of canopy structure and productivity. Notably, Open architecture, which is generally undesirable due to poor canopy structure and reduced field efficiency, showed strong negative correlations with more favorable plant forms. This antagonistic relationship implies that selection against open types may indirectly promote preferred architectures, such as Umbrella. Leveraging such correlated responses through index-based or phenotypic selection provides a practical route for incorporating complex architectural ideotypes into breeding pipelines.

### 3 Genetic architecture of BranchlevelNum and candidate gene discovery

The convergence of BLINK and MLM on a shared locus on chromosome 2 provides strong evidence that this region represents a robust and biologically meaningful determinant of variation in BranchlevelNum. Although BLINK detected multiple significant associations across the genome, its strongest signal coincided with the same chromosome 2 region identified by the more conservative MLM, indicating that this locus is resilient to differences in model assumptions and statistical power. The spatial concentration of associated markers within this chromosome 2 region, together with their concordant effects across analytical models, is consistent with the presence of a single, major genetic factor influencing BranchlevelNum.

Our findings corroborate earlier linkage mapping reports, which located major QTLs for plant height and first branch height on cassava chromosome 2 highlighting its central role in controlling shoot architectural components [47]. While previous studies have frequently associated architectural traits such as branching level and on chromosomes 1, 5, 8, 11, 13, 14, 15, and 18 [12,32,48], the detection of a stable and specific locus on chromosome 2 in the present study provides new insight into additional genomic regions contributing to shoot morphology. The specificity of this signal to BranchlevelNum further supports the feasibility of incorporating branching traits into ideotype-oriented breeding strategies aimed at improving reproductive synchronization and agronomic efficiency.

Functional annotation of genes within the chromosome 2 QTL region revealed several candidates with compelling roles in plant developmental regulation. Notably, this region contains genes associated with cell cycle regulation, organogenesis, and meristem function, including a homolog of SCARECROW-LIKE 28, suggesting that variation in branching level may be governed by differences in axillary meristem activity and local organogenic competence [40,49]. Such genes are known to play key roles in defining meristem identity and regulating lateral bud fate in model systems[2,8]. In addition, the presence of genes related to reproductive development, such as Hapless 2, indicates potential coordination between vegetative and reproductive processes in cassava. This is consistent with the fact that branching is required for flowering in cassava and that increased BranchlevelNum is associated with enhanced floral output [14,50]. Finally, the identification of multiple cytochrome P450 genes within the QTL region further supports the involvement of hormone-mediated signaling pathways, as these enzymes participate in the biosynthesis and catabolism of auxin, cytokinin, and strigolactones, key regulators of shoot branching and apical dominance [3,7]. These results suggest that variation in cassava branching architecture may involve genetic and hormonal regulators acting within this genomic region.

### 4 Genomic prediction and integration strategy for architecture traits

The genomic prediction analysis conducted in this study provides a strong foundation for the strategic integration of specific architecture traits into cassava breeding programs. Among all traits evaluated, BranchlevelNum emerged as the most predictive, achieving a genomic prediction accuracy of 0.44. This level of performance is comparable to the highest predictive abilities reported for key agronomic traits in cassava, including root yield and disease resistance, which typically range from 0.40 to 0.53 in African germplasm [18,31,37,51]. The relatively high predictability of BranchlevelNum aligns with its robust GWAS signal and high broad-sense heritability, reinforcing its value as a reliable target for early-generation genomic selection. Beyond its genetic tractability, this trait holds practical breeding relevance due to its established association with flowering phenology and canopy architecture, both of which are critical for hybridization success and field performance.

PlantHeight, FirstBranchHeight, and Open type exhibited moderate genomic prediction accuracies (approximately 0.25–0.26), suggesting that these traits are best leveraged through index-based selection or stage-specific culling, rather than as primary early-generation genomic targets. In practice, selection against extreme values for these traits may be most effective during advanced testing stages or under controlled environments where environmental noise is reduced. In contrast, Compact and Umbrella plant types showed limited direct predictability; however, their strong relevance to agronomic performance and reproductive efficiency supports their use as qualitative ideotype constraints in breeding decisions. These plant forms are favored by breeders and farmers due to advantages in harvestability, lodging resistance, canopy efficiency, flowering synchrony, and crossing efficiency. Accordingly, rather than relying on direct genomic prediction, Compact and Umbrella architectures may be most effectively incorporated through phenotypic screening, indirect selection via correlated traits, or penalty-based exclusion of undesired open architectures within selection indices. This combined strategy allows breeders to align genomic selection with practical ideotype goals while maintaining flexibility in trait prioritization.

The comparison between additive and additive-dominance genomic prediction models revealed no substantial gain from including dominance effects. This indicates that additive genetic variation captures most of the heritable signal for the architectural traits examined, simplifying model deployment and enabling the use of additive GEBVs in standard breeding operations without loss of predictive performance.

Across architectural traits, clear differences emerged in genetic stability and predictability, indicating that not all traits are equally suited for selection at the same stage of the breeding pipeline. Traits such as BranchlevelNum and Cylindrical plant type combined relatively high heritability, robust GWAS signals, and superior genomic prediction accuracy, suggesting that they capture stable genetic variation that can be reliably exploited early in selection. In contrast, traits including PlantHeight, FirstBranchHeight, and plant shape categories were more strongly influenced by environmental variation or qualitative assessment, indicating that their optimization may benefit from later-stage evaluation or indirect selection. Together, these patterns indicate that architectural traits differ in their timing and mode of effective selection, rather than responding uniformly to a single breeding approach.

## Conclusion

This study provided the first large-scale comparative analysis of plant architectural traits across Nigerian cassava breeding programs, integrating phenotypic trends, heritability estimates, trait correlations, genome-wide association analyses, and genomic prediction. Together, these approaches reveal that cassava architecture is under substantial genetic control and can be modified through selection without compromising yield performance. Although NRCRI and IITA progress through their breeding pipelines with distinct architectural trajectories, both programs ultimately advance varieties with similar release-stage plant forms, indicating convergence toward shared agronomic and farmer-preferred ideotypes.

Among the traits evaluated, BranchlevelNum and Cylindrical type emerged as particularly valuable targets due to their high heritability, genetic stability, and limited trade-offs with yield-related traits. The identification of a major-effect locus on chromosome 2, supported by both BLINK and MLM models, and the presence of functionally relevant candidate genes involved in meristem regulation, organ development, and reproductive processes, provide strong evidence that branching architecture is genetically coupled with reproductive competence in cassava. These findings establish BranchlevelNum as a robust and genetically tractable trait with direct relevance for hybridization efficiency and breeding system performance.

Importantly, this work bridges biological insight and applied breeding by demonstrating that architectural traits can be explicitly incorporated into selection strategies using genomic tools. A tiered breeding framework is supported, in which highly heritable and predictive traits such as BranchlevelNum and Cylindrical type are prioritized for early-generation genomic selection, while more environmentally responsive traits are refined in later stages or specific production contexts. Although Compact and Umbrella architectures exhibited lower direct predictability, their strong relationships with other architectural traits and their importance as breeder– and farmer-preferred ideotypes indicate that they remain relevant targets through indirect or index-based selection.

Looking forward, the integration of multi-trait genomic prediction models that jointly consider architecture, yield, and reproductive traits will be essential for accelerating genetic gain in cassava. By aligning molecular evidence with agronomic performance and stakeholder preferences, breeding programs can more effectively develop cassava varieties that are structurally optimized, flowering-efficient, and well suited to diverse agroecological and production systems.

## Supporting information

**S1 Fig: Q-Q Plots of Standardized Residuals for PlantHeight, FirstBranchHeight and BranchlevelNum across breeding stages in Nigeria Cassava germplasm**

**S2 Fig: Scree plot showing the percentage of explained variance by the principal components from the SNP marker matrix analysis**

**S3 Fig: Manhattan plots of the of the Bayesian-information and Linkage-disequilibrium Iteratively Nested Keyway (BLINK) model and the mixed linear model (MLM) results for plant height architecture traits across 18 chromosomes in 3,113 cassava accessions from the IITA and NRCRI breeding programs.**

**S1 Table: ANOVA Model and Summary of Results for PlantHeight, FirstBranchHeight and BranchlevelNum**

**S2 Table: Pairwise linkage disequilibrium (r²) estimates among significant SNPs within the chromosome 2 QTL for *BranchlevelNum***

**S3 Table: Exploration of Candidate Genes Linked to Number of Branching Levels through functional annotation in Cassava (according to Phytozome v13 and NCBI)**

## Acknowledgments

The authors gratefully acknowledge the Bill & Melinda Gates Foundation for financial support. We thank both the International Institute of Tropical Agriculture (IITA) and the National Root Crops Research Institute (NRCRI) for their essential contributions to field trial implementation and data collection, as well as for providing access to cassava germplasm, multi-location trials, and phenotypic datasets. We also acknowledge the NextGen Cassava project and its partners for access to genotypic data and technical support.

## Data availability statement

The original contributions presented in the study are included in the article/Supplementary Material. All data files and analysis scripts are available online on the github url address https://github.com/Opamelas83/Plantarchitect

## Funding

The author(s) declare that financial support was received for the research and/or publication of this article. This work was supported by the Bill & Melinda Gates Foundation through the “Next Generation Cassava Breeding project” (https://www.nextgencassava.org; grant no. INV-007637) managed by Cornell University.

## Notes

### Competing Interest Statement

The authors have declared no competing interest.

